# Rat bone marrow derived mesenchymal stem cells differentiate to germ cell like cells

**DOI:** 10.1101/418962

**Authors:** Kuldeep Kumar, Kinsuk Das, AP Madhusoodan, Ajay Kumar, Purnima Singh, Tanmay Mondal, Sadhan Bag

## Abstract

The *in vitro* differentiation of rMSCs provides an excellent model for studying cell commitment and their potential in stem cell technology. We have demonstrated that rat bone marrow derived MSCs are able to differentiate into germ-line cells *in vitro* which has an enormous scope in the advancement of fertility research. In future, this technique can be utilized in different domestic animal species for propagation of elite germ plasm.

**Abstract:** Germ cells undergo distinctive male or female pathways to produce spermatozoa or oocyte respectively essential for sexual reproduction. Mesenchymal stem cells (MSCs) have the capacity of trans-differentiation to form the multiple lineages of cells of mesoderm, endoderm, and ectoderm origin. Herein, MSCs were isolated from rat bone marrow and characterized by their morphological features, expression of surface markers by immunophenotyping and molecular biology tools as well as self renewal and differentiation capability. Thereafter, by inducing these cells with retinoic acid we could able to show that bone marrow derived MSCs are able to trans-differentiate into male germ cell-like cells which were further characterized by the expressions of germ cell specific markers. This *in vitro* study for the generation of germ-like cells suggests that bone marrow MSCs can be a potential source of germ cells that could be a sustainable source of sperm / oocyte production for potential therapeutic applications in future. Moreover, this technique can be applied in different domestic animal species for propagation of elite germ plasm.

## Introduction

Bone marrow (BM) contains two types of cells; hematopoietic stem cells are responsible for the production of blood cells and, mesenchymal stem cells (MSCs) which have differentiation capability into the various cell lineages including adipocytes, chondrocytes and osteoblast (Friedenstein et al., 1974; Hunt et al., 2987; Prockop et al., 1997). Initially these cells were named as plastic-adherent or colony-forming unit fibroblast (CFU-F) which later on known as bone marrow stromal cells, mesenchymal stem cells (MSC) due to their heterogenous character and potentiality to differentiate into the various cells phenotypes (Jiang et al., 2002; Pittenger et al., 1999).

During last decade, MSCs have been isolated not only from bone marrow but also from different tissues of diverse species such as mice, rat, human, caprine, canine, equine, bovine, porcine, guinea pig etc. which could be differentiated into numerous cell types of mesodermal and non-mesodermal origin under appropriate conditions (Sung et al., 2008; Lotfy et al., 2014; Francis et al., 2009; Kumar et al., 2013; Das et al., 2017; Lange-Consiglio et al., 2013; Raoufi et al., 2011; Chen et al., 2016; Aliborzi et al., 2016). Few research works have demonstrated that MSCs not only inherently show a number of germ cell (GCs) characteristics but also in presence of certain chemical cue it may be propelled to GCs differentiation (Nayernia et al., 2006; Drusenheimer et al., 2007; Huang et al., 2010; Shirazi et al., 2012; Mazaheri et al., 2011). This differentiation potentiality into germ cells has paved the way for capitalizing these cells as a potential solution for infertility. Studies ranging from the chemical induction of MSCs to transplantation of MSCs into gonads have been performed to produce GCs (Nayernia et al., 2006). In most of the *ex vivo* studies, retinoic acid (RA) have been used as potential cue for differentiation of MSC into germ cells because of its important regulatory role in embryonic patterning and development (Nayernia et al., 2006; Drusenheimer et al., 2007; Huang et al., 2010; Koubova et al., 2006).

MSC derivation from rat bone marrow has been reported earlier but, to the best of our knowledge its differentiation capability towards germ cell-like cells has not yet been recorded. Aim of the present study was to isolate MSCs from rat bone marrow, and to explore their differentiation potentiality towards germ cell-like cells. We successfully isolated and characterize MSCs and, were made to germ cell (GC) differentiation *ex vivo* with the induction of retinoic acid (RA) for 21 days. Confirmation of GCs was completed by morphological changes in MSCs over the time, germ cell specific gene expression by reverse transcription polymerase chain reaction (RT-PCR), immunophenotyping of cells with GC markers followed by flow cytometry based quantification of differentiated cells. The findings of this study may possibly be an insight cue for its prospective application in reproductive health.

## Results

### Isolation and characterization of bone marrow derived rMSCs

Rat bone marrow derived cells were noticed attached and proliferated onto the polysterin coated plastic surface of culture flask within 3 days after seeding. Initially different morphology of round and spindle shape of these cells was visualized which on subsequent passage falttened further to became fibroblastic in shape. Expanded cells after four passage appeared morphologically homogenous (Fig 1A).

**Figure. 1:**
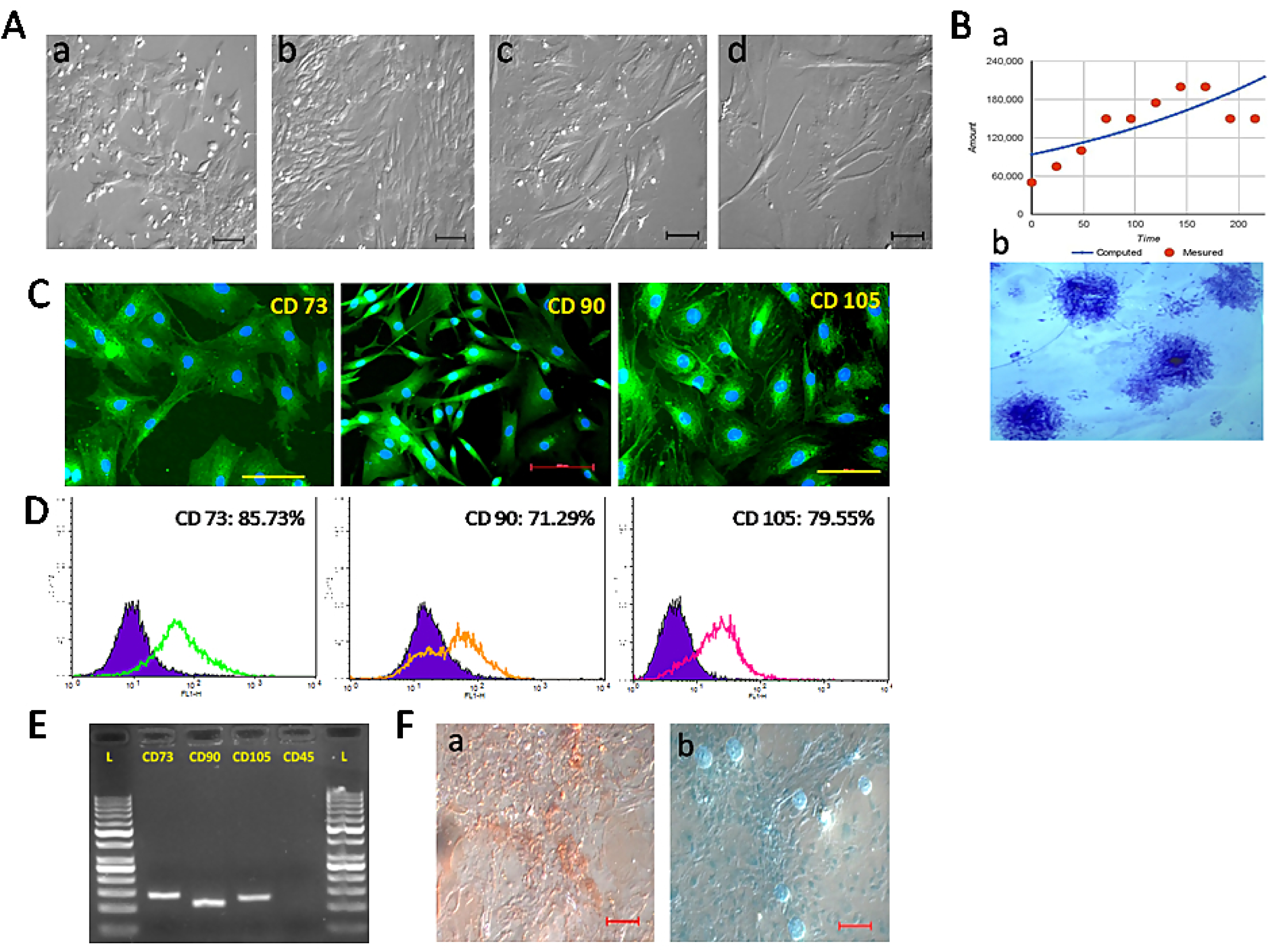
(A-F) Characterization of rMSCs. (A) Morphology of rMSC at different passage (a)P 0, (b) P1, (c) P4 and (d) P6. (B) (a) Growth curve and (b) CFU assay of rMSC (C). Immunofluorescence staining of MSC specific surface markers CD 73, 90 and 105 (D) Values represent the mean percentage of positively stained cells as analyzed by flow cytometry (E) Expression of MSC specific genes assessed by RT-PCR (F) Differentiation potential of rMSCs into mesodermal lineages. (a) osteocytes (alizarin red staining), (b) chondrocytes (alcian blue staining). Scale bar = 100 μm

Growth curve of rMSCs was obtained by counting the cells manually and the average of population doubling time was calculated based on the logarithmic growth phase. The average population doubling time was recorded 187.32h. CFU assay provided an evidence of these cells’ proliferative and clonogenic capacity in culture system (Fig.1B).

In immunocytochemical staining cells were positive for MSCs specific surface markers, CD 73, CD 90, and CD 105 (Fig. 1C). Fluorescence activated cell sorting (FACS) revealed that the cells were positive for CD 73 (85.73%), CD 90 (71.29%) and CD 105 (79.55%) (Fig. 1D). Further, gene expression followed by gel electrophoresis study revealed that distinct bands were expressed for MSC specific genes CD73, CD90, and CD105 whereas no band was noticed for CD45 (Fig. 1E).

Cells were able to differentiate to osteocytes, and chondrocytes under standard *in vitro* differentiating conditions. After osteogenic differentiation for 21 days prominent mineralized nodules were visualized in alizarin staining. Proteoglycan accumulation was noticed by Alcian Blue staining after 21 days of chondrogenic differentiation (Fig. 1F).

### *In vitro* differentiation of rMSCs into germ cell like cells

After induction we noticed the effects of retinoic acid (RA) treatment on rMSC in terms of morphological changes at various time points. At the end of the treatment period changes in some of the cells’ morphology towards germ-like cell in the culture was visualized (Fig. 2). Immunocytochemical staining for the germ cells markers like stella and fragilis were found positive in treated cells after 21days (Fig. 3A). These positive cells were quantified by FACS analysis, revealed that RA treatment led to the generation of 56.75% and 39.51% stella and fragilis positive cells, respectively (Fig. 3B). In RA treated cells, RT-PCR results detected the expression of stella and fragilis genes which are believed to be expressed at early stage of germ cell development. Moreover, we noticed the expression of other germ cell marker genes like c-Kit, Stra8, DAZL, Tex18, INGA6 and TP2 (Fig. 3C, D).

**Figure 2:**
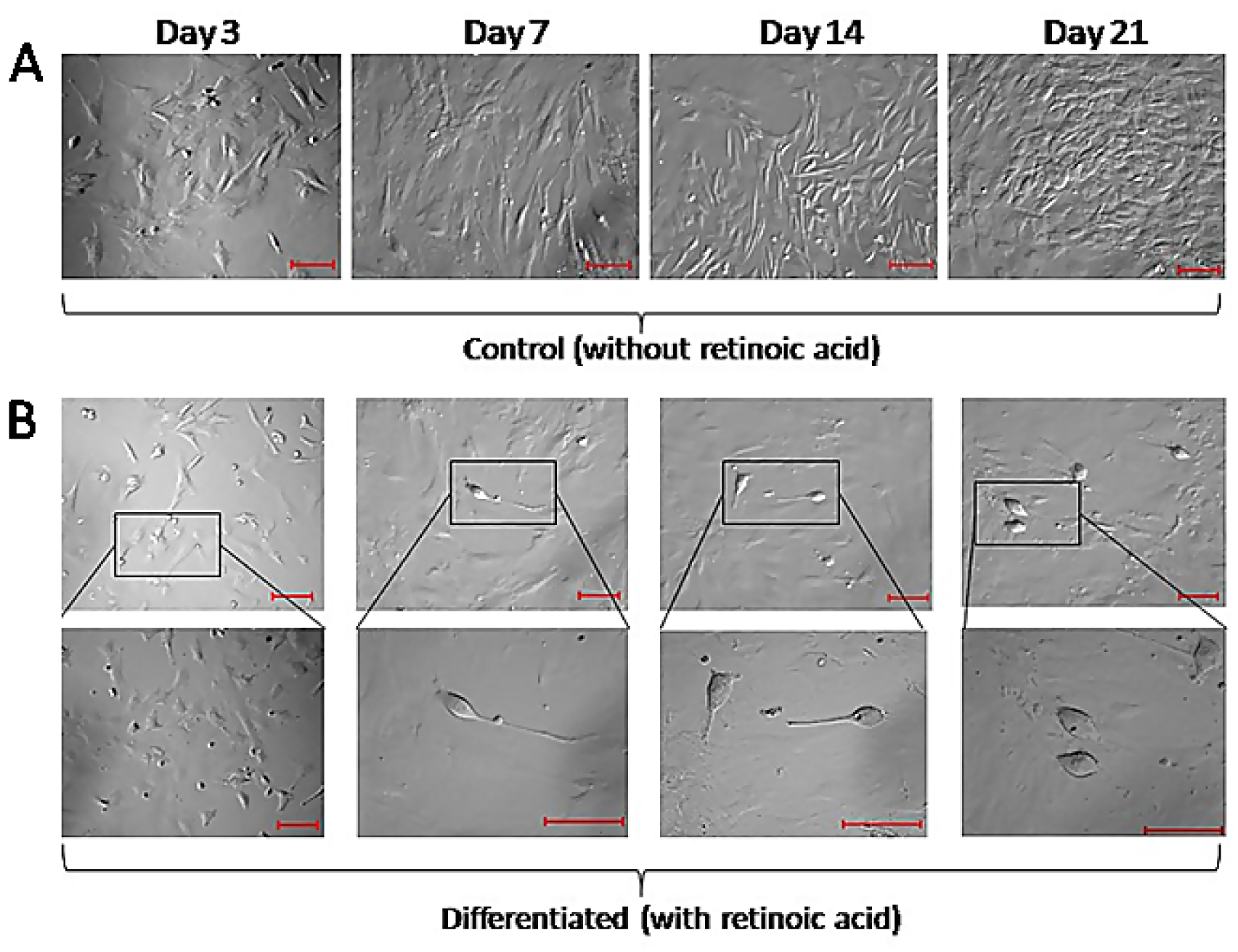
*In Vitro* Differentiation of rMSC into germ cell-like cells on different days. (A) Control undifferentiated (B) Differentiated with RA supplementation. Scale bar = 100 μm.

**Figure 3:**
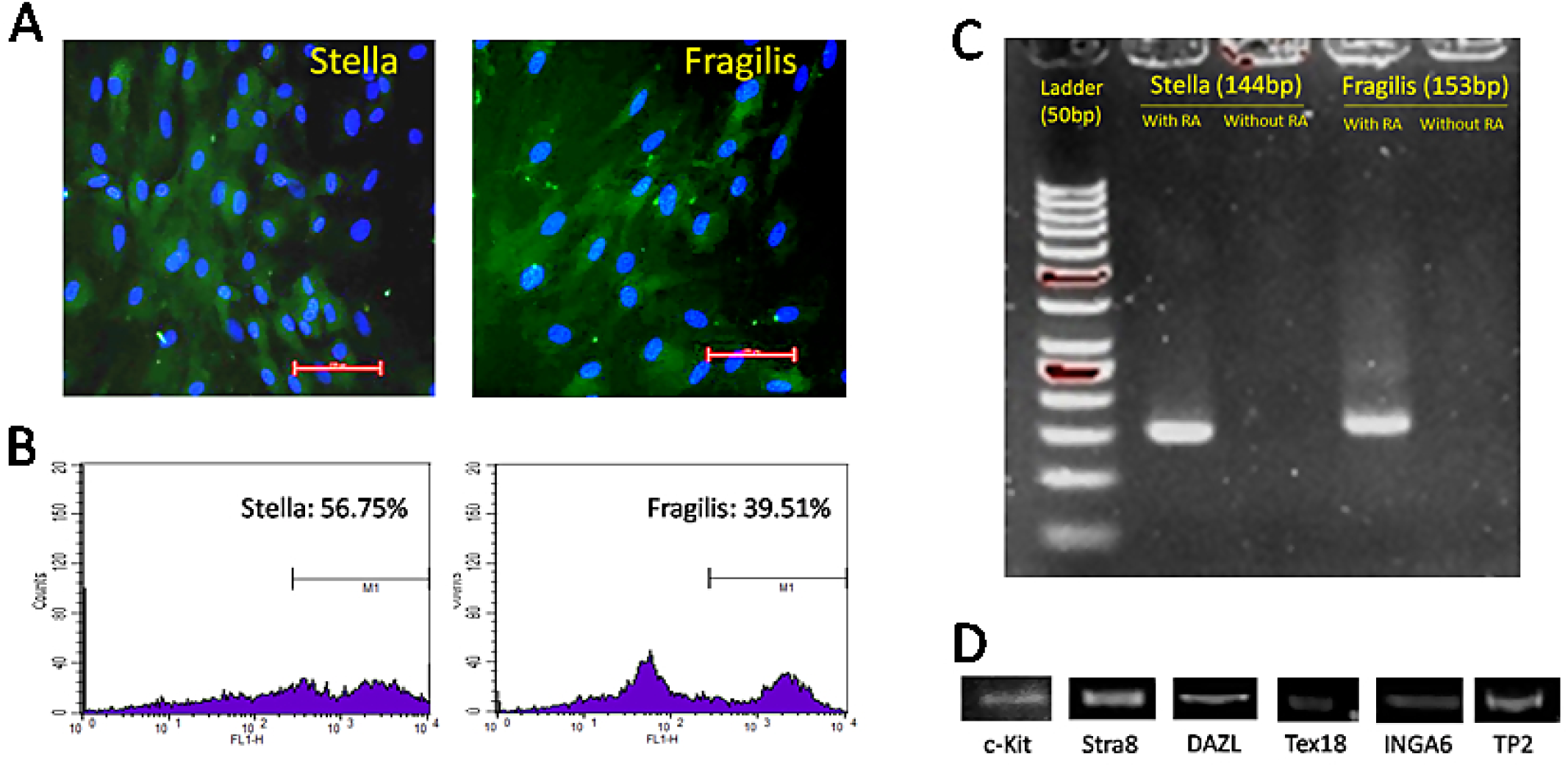
*In Vitro* characterization of germ cell-like cells. (A) Immunofluorescence staining after treatment with RA for 21 days (Scale bar 100μm) (B) FACS analyses of rMSCs derived germ cells, labeled with FITC conjugated antibodies against Stella (56.75%) and Fragilis (39.51%) (C) Molecular characterization of rMSC differentiated cells into germ cells by Stella and Fragilis (D) Molecular study of spermatogonial stem cells markers - c-Kit, Stra8, Dazl, tex18, Inga6 and TP2

## Discussion

MSCs are a group of cells that can be found in various organs and tissues such as the bone marrow (BM), amniotic fluid, lungs, fat, peripheral blood, dental pulp, spleen, muscles, kidneys, and umbilical cord blood etc. Under appropriate conditions, they have the capacity to differentiate into numerous cell types and good option for use in cell therapy. Therefore, infertility researchers are motivated to use these cells as a potential solution for infertility. The approach of oocytes/ova production from MSCs may significantly improve fertility in female, elite animals and conservation of the germplasm of endangered and extinct species in future. The germ cell generation from MSC of various sources has enormous potential to increase animal reproduction. It offers the ability to produce animals of elite genetic potential from germ cells generated from MSC, if the ovum and semen are not available because of non-genetic reasons.

Isolation and purification of rat MSCs is more challenging than other species due to its low abundance in bone marrow and also due to unwanted growth of non-mesenchymal cells during *in vitro* cell culture (Eslaminejad et al., 2006). In the present study, spindle-shaped fibroblast-like cells from rat bone marrow were purified by repeated medium change at initial phases of culture. Thereafter, in subsequent passage cells displayed the characteristic spindle shaped morphology of MSC which is commonly found in rat (Meirelles et al., 2003; Quiroz et al., 2008; Niyaz et al., 2012). The isolated cells displayed the formation of clones from single cells as determined by CFU assay. This assay demonstrate the capacity of cell to generate clones i.e. self-renewal ability, which is a typical characteristic of stem cell populations. Population doubling time was performed for measuring the time taken by the cells in doubling their number; result indicated that rat bone marrow MSCs are highly proliferative in nature. Lotfy et al., (2014) reported a higher proliferation rate of rat MSCs when derived from bone marrow as compared to its adipose tissue origin. Isolated cells expressed specific mesenchymal stem cell markers positive for CD73, CD90, and CD-105 whereas, negative for CD-45. Apart from this, the cells were able to differentiate into osteogenic and chondrogenic lineages. Similar reports have been published by other researchers while characterizing rMSC (Asumda et al., 2011; Sun et al., 2011; Zangi et al., 2006; Davies et al., 2015). Therefore, we presume that cell population isolated from rat bone marrow is of rMSCs and, these cells were used for our subsequent experiments.

The embryonic stem (ES) cells can be differentiated into germ cells in either by attachment culture or from embryoid bodies *in vitro* (Tilgner et al., 2008; Yamauchi et al., 2009; Lacham et al., 2006). Germ cells like cells have also been generated spontaneously from human ES cells which expressed different pre-meiotic and post-meiotic germ cells marker along with stem cells markers (Clark et al., 2006; Ginsburg et al., 1990). However, only few studies envisaged the potential of mice MSC towards germ cell lineage differentiation. These studies reported that mice MSCs under appropriate *in vitro* conditions can generate GCs (Nayernia et al., 2006; Drusenheimer et al., 2007; Huang et al., 2010; Shirazi et al., 2012). In the present study, we could able to generate germ cell-like cells after incubating the rMSC in RA following the method reported by Nayernia et al., (2006) in mice. RA is a crucial signaling molecule during vertebrate development, plays a key role in cell differentiation, proliferation and apoptosis (Miano et al., 2001). Moreover, it is considered as one of the important factors for inducing meiosis in differentiating mouse germ cells (Bowles et al., 2007). RA at high concentration (10^-5^ to 10^-6^ M) induces the formation of germ cells while its low concentration (10^-8^ to 10^-9^ M) promotes differentiation of MSCs towards smooth muscle and myocardial cells (Geijsen et al., 2004; Silva et al., 2009; Wobus et al., 2002). Herein, the differentiated cells resembled to the primordial germ cells of mice, human and primate (Shirazi et al., 2012).

Stella is a germ cell specific gene and amongst the first genes express in the differentiation process of GCs. It is believed that Stella is highly expressed in adult testicular spermatogonia (Lacham-Kaplan 2004). Moreover, it is involved in triggering GC competence and specification and in the delimitation of primordial GCs from their surrounding somatic cells (Aflatoonian and Moore 2005; Hayashi et al. 2007; Saitou et al. 2002; Saitou 2009). Expression of this marker in differentiated MSCs has been documented earlier (Nayernia et al. 2006; Qiu et al. 2013). Fragilis or interferon-induced transmembrane protein 3 (ifitm3) is also a GC-specific marker and an important factor for initiation of GCs specification and competence (Lacham-Kaplan 2004; Lange et al. 2003). In our findings, we mostly focused on these two markers and, by immunophenotyping and gene expression studies and, could able to establish the differentiation of rat MSCs towards germ cell-like cells.

We also tested the differentiated cells for the expression of different germ cells specific markers viz. c-kit, stra8, DAZL, tex18, INGA6, TP2 by RT-PCR analysis and noticed the positive expression. Previous studies also reported that these markers are expressed during GC generation from MSCs under appropriate *in vitro* conditions (Saitou et al., 2002; Wang et al., 2001). DAZL contributes to the preliminary primodial germ cell formation by limiting both pluripotency and somatic differentiation (Niu et al.2014; Chen et al., 2014), and continues its expression in various types of goniocysts in prenatal and postnatal testes (Brekhman et al., 2000). In adult mouse and human testes DAZL is detected in spermatogonia, spermatocytes, and early round spermatids. Firstly it appears in mitotic spermatogonia, reaches peak in the cytoplasm of pachytene spermatocytes and ended in late sperm formation stage (Niu et al.2014). Previous studies also reported the ectopic expression of DAZL successfully promotes meiotic progression of PGC-like cells in human ESCs and iPSCs, as well as in DAZL-derived mESCs (Medrano et al., 2012; Ramos-Ibeas et al., 2015). Stra8 has emerged as a gatekeeper for the entry of germ cells into meiosis which is indispensable for premeiotic DNA replication and subsequent entry into the prophase of meiosis I (Mark et al., 2008; Baltus et al., 2006). Stra8 was previously used to construct the germ cell specific reporting vector Stra8-EGFP in the generation of male germ cells from mouse iPSCs and spermatogonial stem cells in vitro (Li et al., 2014). c-Kit has a similar effect and essential to the process of germ cell migration, proliferation, and survival in the gonadal anlage, and it is also involved in the reprogramming of PGCs (Hoyer et al, 2005). The molecular markers such as fragilis, stella, c-Kit, Tex18, Stra8, Dazl, TP2 and inga6 analyzed in these study have been identified earlier as the evidence of germ cell differentiation and for spermatogonia and SSCs (Saitou et al., 2002; Vincent et al., 1998; Wang et al., 2001; Oulad-Abdelghani et al., 1996; Cooke et al., 1996 and Shinohara et al., 1999).

## Materials and methods

### Experimental Animals

The adult Sprague dawley rats were collected from the laboratory animal house, Indian Veterinary Research Institute, Izatnagar, Bareilly and maintained under standard management practice. The animal care, handling as well as experimental protocols were approved by the Institutional Animal Ethics Committee (IAEC) of the Indian Veterinary Research Institute, which has the approval of the Committee for the Purpose of Control and Supervision of Experiments on Animals (CPCSEA) under the Ministry of Environment, Forests and Climate Change, Government of India.

### Isolation and culture of rat bone marrow derived MSCs

The isolation of rat MSC was following the protocol described by Nayernia et al., 2006. Briefly, adult rats were sacrificed under anesthesia and, femora and tibias of both the sites were aseptically removed. These bones after thoroughly washing with sterile PBS were flushed out with Dulbecco’s Modified Eagle’s Medium (DMEM) to get the bone marrow cells. The released cells were collected into 90 mm culture dish and finally plated at a density of 10^5^ cells/cm^2^ in tissue culture flask (Nunc, Germany) with DMEM-low glucose medium supplemented with 10% FBS, L-glutamine (2mM), penicillin (100U/mL), streptomycin (100µg/mL) and amphotericin B (0.25µg/mL) (all from Thermofisher), and maintained at 37°C in a humidified atmosphere of 5% CO_2_. Cells upon reaching 70-80% confluence were serially passaged by detaching with 0.25% Trypsin-EDTA (Invitrogen). To define these cell population as multipotent MSCs we characterized them to meet the minimal criteria like morphology, plastic adherent property, colony forming ability, expression of MSC specific surface markers (+/CD 73, CD 90, CD 105 and -/ for CD 45) and the ability for multiple-lineage differentiation *in vitro*. The population of rat MSCs (rMSCs) was expended upto 6 passages for our experiments.

### Growth kinetics and colony forming unit assays of rMSCs

The growth kinetic study was performed by harvesting followed by counting the trypsinized cells manually. rMSCs were cultured in 12 well plate at a plating density of 50,000 cells/well. The number of cells of each plate was counted on every day. The population doubling time of cells was estimated on logarithmic growth phase based of the cell growth curve. To assess the capacity and efficiency for self renewal, cells (Passage 2) were seeded at low density in 6-well culture plate (50 cells/cm^2^) and new fibroblast colonies derived from single cells were counted. After 15 days of culture the plates were fixed and stained with 1% crystal-violet in 100% methanol (Kumar et al., 2016)

### Immunofluorescence staining and flow cytometry of rMSCs

The immunocytochemical staining of rMSCs was done with MSC specific positive markers like CD73, CD90 and CD105 and, with CD45 as a negative marker. The rMSCs were grown over cover slip and, were fixed with 4% paraformaldehyde, washed with PBS, permeabilized with 0.25% Triton X-100 in PBS followed by blocking nonspecific binding sites with 10% normal goat serum in PBS for 1h at RT. The cells were then incubated with primary antibodies (1:100; Santa Cruz) of CD73, CD90, CD105 and CD45 for overnight at 4°C. After thorough washing, the cells were further incubated with FITC conjugated secondary antibodies (1:500; Santa Cruz) for 4 h at darkness followed by counter staining with DAPI ProLong Gold antifade solution (Invitrogen). Cells were imaged under an inverted fluorescence microscope (Carl Zeiss) with Axio Vision 4.0 image analysis system. To evaluate the homogenecity of rat MSCs, cells were analyzed by flow cytometry. Aliquots containing 1 × 10^6^ cells for each marker were separately fixed, permeabilized, blocked followed by incubation with primary and secondary antibodies as per the standard immunocytochemical staining protocol with the same combinations of antibodies used for each marker. Flow cytometer (FACS Calibur, BD Bioscience, USA) settings were established using unstained cells. Data was analysed by recording 10,000 events with Cell Quest Pro software (BD Bioscience, USA).

### In vitro differentiation of rMSCs

To induce osteogenic differentiation, rMSCs were plated in a six well plate tissue culture dish in MSC medium. After 48 h of culture, the media was replaced with osteogenic differentiation medium (DMEM containing 10% FBS and 10 nmol-Dexamethasone, 10 mmol – β-glycerophosphate, 0.3 mM –L-ascorbic acid) upto 21 days with adding fresh differentiation media at every 48-72 h interval. The osteogenic differentiation was visualized by Alizarin red cytochemical staining. For chondrogenic differentiation, the cells were cultured in chondrogenic induction medium consisting of DMEM containing 10% FBS, 100 nmol-Dexamethasone, 1mmol sodium pyruvate, L-ascorbic acid (50µg/ml), TGF-β1 (10ng/ml and ITS 1%). The induction medium was changed after every 3^rd^ day. At the end of the induction, the cells were fixed with 4% paraformaldehyde for 10 min and were stained for the alcian blue staining.

### Gene expression study

Cells were harvested from monolayer culture and washed with PBS. The total RNA was isolated by Quick-RNATM MicroPrep (Zymo Research). Total RNA was quantified, and quality was ascertained (OD 260/280 > 1.6) using NanoDrop (Eppendrof, USA) as well as gel electrophoresis. A total of 1µg RNA was reverse-transcribed to synthesize complimentary DNA (cDNA) using iScript™ cDNA Synthesis kit (Bio-Rad, USA). The expression of MSC specific genes CD73, CD90, CD105 and CD45 was assessed by RT-PCR method with specific primers (Table 1) by using EvaGreen supermix (Bio-Rad). Each PCR product was size-fractionated by 2% agarose gel electrophoresis, and the bands were visualized with a UV trans-illuminator (Bio-Rad).

**Table 1:**
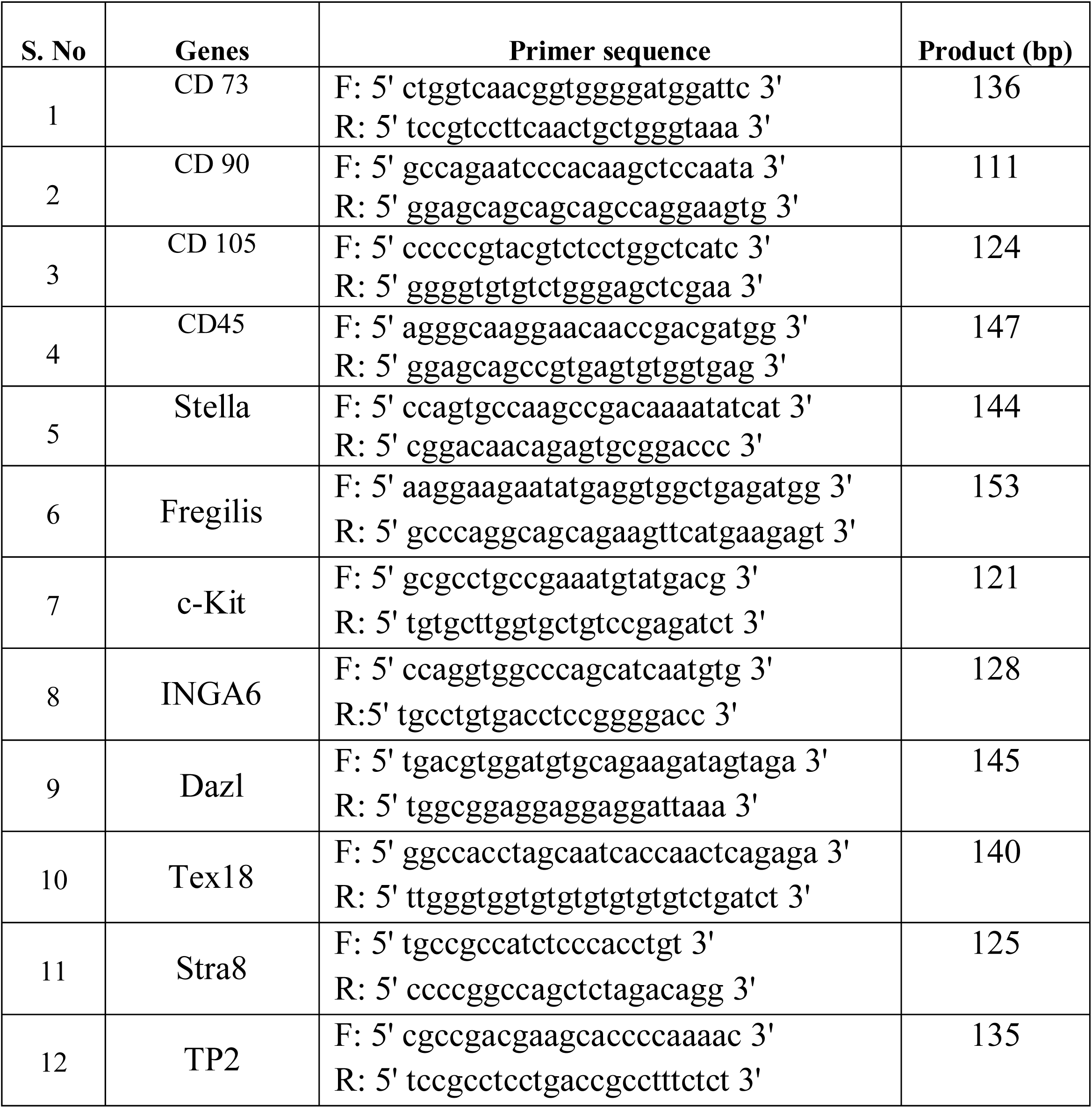
List of primers used in the study

### Induction for germ cell differentiation

The differentiation of rat MSC into GCs was induced by the method described by Nayernia et al., (2006). Briefly, the semi confluent cells were treated in MSC culture medium with supplementation of 10^-6^M RA (Sigma) and maintained at 37°C in a humidified atmosphere of 5% CO2 for 21 days. After induction we looked into the effects of retinoic acid (RA) treatment on rMSC in terms of morphological changes at various time periods of 3, 7, 14 and 21 days as well as expressions of GC-specific genes, immunocytochemical staining followed by flow cytometry at the end.

### Molecular characterization

Total RNA was extracted and cDNA was prepared from the differentiated cells after 21 days. RT-PCR gene expression analysis was carried out using the Real-Time PCR System (Bio-Rad) with rat specific primers (Table 1) for the germ cell marker genes like Stella, Fragilis, c-Kit, Stra8, DAZL, Tex18, INGA6 and TP2. Size-fractionation of PCR products were done as per previously mentioned protocols.

### Immunocytochemistry

In order to confirm the differentiation of rMSCs into GCs, we performed immunostaining for expression of GCs specific markers Stella and Fregilis. rMSCs were differentiated over coverslips which was kept inside each well of the 6 well plate. The cells were subsequently fixed and stained with antibodies according to a previously described method (Kumar et al., 2014)

### Flow cytometry assay of germ cells

Following differentiation, the cells were harvested by trypsinization and collected by centrifugation followed by washing with PBS. Cells for each marker were separately fixed, permeabilized, blocked followed by incubation with primary and secondary antibodies (Stella & Fragilis) as per the standard staining protocol as described for immunocytochemical staining. Flow cytometer (FACS Calibur, BD Bioscience, USA) settings were established using unstained cells. Data was analysed by Cell Quest Pro software (BD Bioscience, USA).

